# Exclusive functional subnetworks of intracortical projections neurons in primary visual cortex

**DOI:** 10.1101/153247

**Authors:** Mean-Hwan Kim, Petr Znamenskiy, Maria Florencia Iacaruso, Thomas D. Mrsic-Flogel

## Abstract

The rules by which neurons in neocortex choose their synaptic partners are not fully understood. In sensory cortex, intermingled neurons encode different attributes of sensory inputs and relay them to different long-range targets. While neurons with similar responses to sensory stimuli make connections preferentially, the relationship between synaptic connectivity within an area and long-range projection target remains unclear. We examined the local connectivity and visual responses of primary visual cortex neurons projecting to anterolateral (AL) and posteromedial (PM) higher visual areas in mice. Although the response properties of layer 2/3 neurons projecting to different targets were often similar, they avoided making connections with each other. Thus, projection target acts independently of response similarity to constrain local synaptic connectivity of cortical neurons. We propose that this segregated connectivity rule reduces the crosstalk between different populations of projection neurons, allowing top-down signal to modulate the activity of these output channels independently.

## Introduction

Uncovering the relationship between the connectivity and function of cortical neurons is fundamental to understanding the circuit mechanisms of information processing. While a neuron’s receptive field, determined by the pattern of inputs it receives, defines a set of stimulus features that drive it to fire, its local and long-range axonal projections define its impact on other neurons. Recent work has begun to uncover the rules by which the receptive fields of excitatory neurons constrain their local connectivity. In layer 2/3, pyramidal neurons preferentially connect to neurons that receive common synaptic input (Yoshimura et al., 2005; Morgenstern et al., 2016) or respond to similar visual stimuli (Ko et al. 2011; Cossell el al. 2015; Lee et al., 2016). However, little is known about how neurons’ long-range projection targets relate to their local connectivity and functional properties.

Paired recordings of layer 5 pyramidal neurons indicate that connectivity between different projection classes is asymmetric (Brown and Hestrin 2009; Kiritani et al. 2012; Morishima et al. 2011). The large pyramidal tract type neurons in layer 5, such as corticospinal cells in motor cortex, and cortico-collicular cells in visual cortex, receive more input than they provide to intratelencephalic (IT) projection neurons (e.g. corticostriatal or corticocortical cells), suggesting they integrate local input before broadcasting information to subcerebral motor-related structures. Whether similar rules govern the local connections of projection neurons targeting different cortical areas is not known.

Signals from primary sensory areas of the neocortex are distributed to downstream areas that work in parallel to deconstruct the sensory scene. Cortical projection neurons innervating different areas may already specialize in encoding different attributes of sensory input (Movshon and Newsome 1996; Sato and Svoboda, 2010; Jarosiewicz et al., 2012; Glickfeld et al. 2013; Chen et al. 2013; Yamashita et al. 2013; Kim et al. 2015; Lur et al., 2016), suggesting they comprise separate output channels sub-serving different sensory and behavioural functions.

In the mouse visual system, signals are relayed from primary visual cortex to a number of higher visual cortical areas (Wang and Burkhalter 2007; Garrett et al., 2014; Zhuang et al., 2017), which differ in their functional properties (Marshel et al. 2011; Andermann et al. 2011; Roth et al., 2012). Among these, the anterolateral visual area (AL) specializes in processing fast moving low spatial frequency stimuli, while the posteromedial visual area (PM) responds primarily to high spatial frequencies. The axonal boutons of V1 neurons in these areas share these biases (Glickfeld et al. 2013; Matsui and Ohki, 2013). To understand how the long-range targets of V1 neurons that comprise these projections relate to their local connectivity, we measured the connection rates between neurons projecting to areas AL and PM using multiple whole-cell patch-clamp recordings. These recordings reveal that layer 2/3 neurons tend to make connections with cells projecting to the same long-range target, while connections between cells projecting to different areas are rare. These observations show that the long-range projection target of cortico-cortical neurons constrains their potential local synaptic partners.

We have previously observed that local connectivity of pyramidal neurons in mouse V1 is correlated with their selectivity for visual stimuli (Ko et al. 2011; Cossell et al. 2015). To examine how the specific connectivity of AL and PM projection neurons relates to their visual response properties, we recorded the activity of these cell populations simultaneously in awake mice using two-photon calcium imaging. Although they differed in their speed tuning and direction selectivity, both populations were highly heterogeneous in their visual responses. The activity patterns of cell pairs projecting to the same area tended to be more similar than of those projecting to different ones. However, these differences were not sufficient to explain the segregated connectivity of these populations, suggesting that other mechanisms play a role in preventing cross-talk between different projection neuron populations.

## Results

### AL and PM projecting neurons are largely non-overlapping populations

To what extent do V1 projections to areas AL and PM originate from distinct populations of neurons? To answer this question, we injected the tracer Choleratoxin B or retrograde Pseudorabies Virus (PRV) (Oyibo et al., 2014) into these areas, identified using intrinsic signal imaging, and examined the distribution of retrogradely labelled neurons in V1 (Figure 1a-d). Cells labelled with both tracers and therefore projecting to both higher visual areas were rare and were found primarily in deep cortical layers (Figure 1d). Therefore, AL and PM projecting neurons in V1 comprise intermingled, non-overlapping populations.

**Figure 1.**
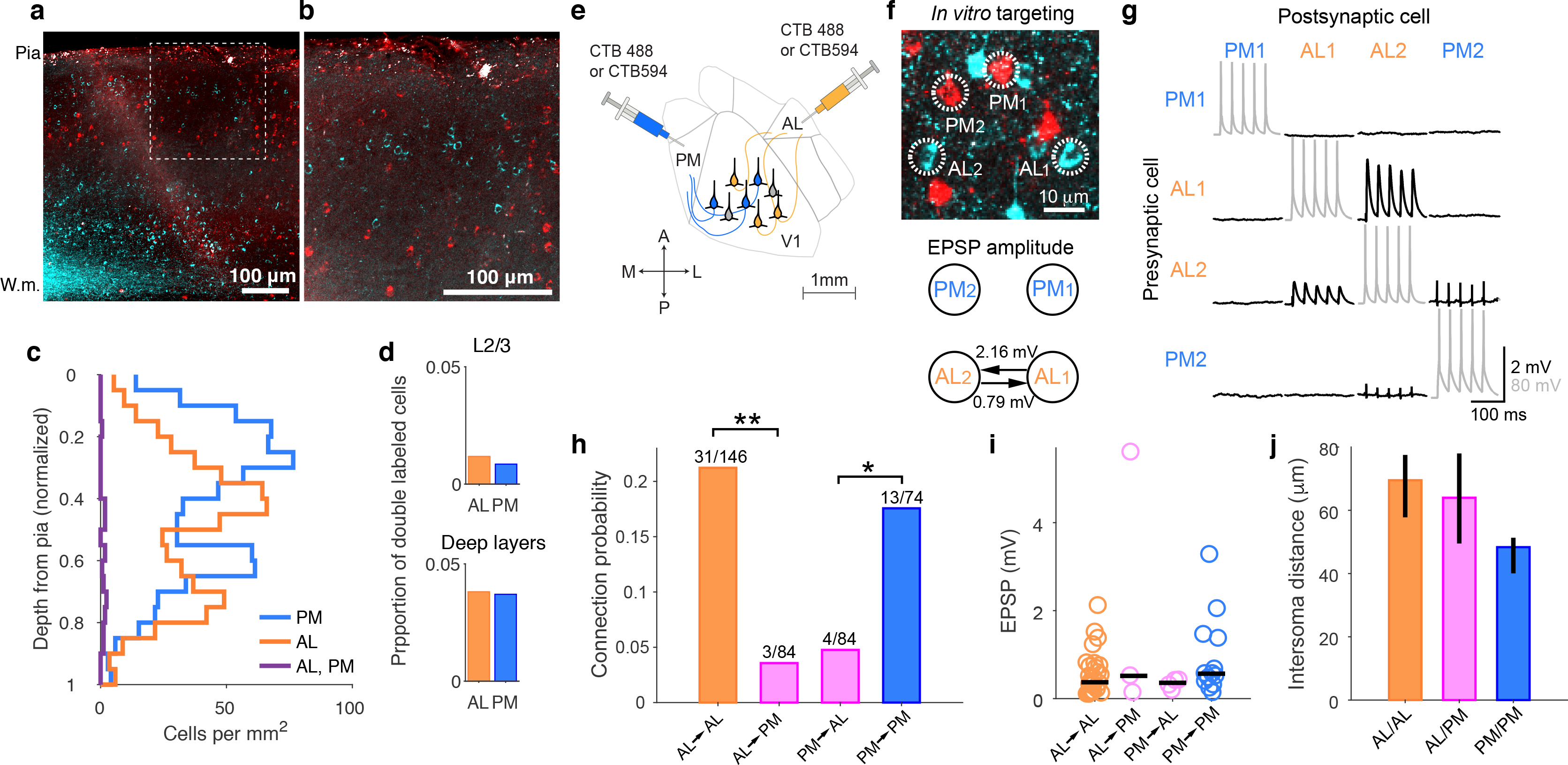
Local recurrent connectivity of AL and PM projection neurons. **a-b**. AL and PM projection neurons labelled by Cholera Toxin B conjugated to AlexaFluor-488 (cyan) and −594 (red) in a coronal section of primary visual cortex. Panel b shows highlighted area of panel **a**. W.m – white matter. **c**. Depth distribution of retrogradely labelled neurons in V1 from 29 sections in 2 mice. **d**. Frequency of double labelled cells as a fraction of the entire AL and PM projection population in L2/3 (upper panel) and deep layers (lower panel) in V1. **e**. Strategy for labelling of AL and PM projection neurons for in vitro recordings. **f**. Example image of CTB labelled AL and PM projection neurons in an *in vitro* brain slice and their connectivity. **g**. An example of membrane potential recordings from four simultaneous recorded cells in current-clamp mode. Two AL projection (AL1, AL2) and two PM projection (PM1, PM2) neurons were targeted, and 5 brief step currents were injected into each cell to evoke action potentials. A reciprocal excitatory connection was found between cells AL1 and AL2. Gray traces indicate presynaptic spikes evoked by current injections, black traces indicate corresponding postsynaptic voltage responses. **h**. Connection probabilities between AL and PM projection neurons. Connection probabilities between neurons projecting to different targets were significantly lower than those projecting to the same target (** < 0.01, * < 0.05 from Fisher’s exact test). **i**. Magnitude of EPSPs between connected AL and PM projection neurons. Black lines indicate medians. **j**. Intersoma distances measured on the tip of patch pipettes after recordings for patched AL and PM projection neuron pairs in slice.

### AL and PM projection neurons form exclusive subnetworks in layer 2/3 of primary visual cortex

Excitatory neurons in layer 2/3 form synaptic connections with only a small fraction of nearby neurons that shares their similar visual selectivity (Ko et al. 2011; Cossell et al. 2015). To understand if the projection target of layer 2/3 neurons in V1 further constrains their local connectivity, we measured the connection rates between different populations of corticocortical projection neurons in layer 2/3. After functionally mapping V1 and surrounding cortical areas using intrinsic signal imaging (see Methods), we injected fluorescent the retrograde tracer Cholera Toxin B (CTB)-conjugated with AlexaFluor-488 and AlexaFluor-594 into areas AL and PM (Figure 1e; see Methods). After allowing at least 3 days for the retrograde transport of the tracers, we assessed the connectivity of retrogradely-labelled AL and PM projection neurons in layer 2/3 of V1. We targeted up to six neurons in acute slices for simultaneous whole-cell patch-clamp recording (Figure 1f-g) to access their synaptic connectivity, strength, and intrinsic membrane properties (Song et al., 2005; Ko et al., 2011; Hofer et al., 2011; Cossell et al., 2015).

The overall connectivity rate of retrogradely-labelled neurons was 13.1 % (51/388), consistent with previous reports for randomly sampled pyramidal neurons in layer 2/3 of mouse neocortex (Ko et al., 2011; Cossell et al., 2015; Lefort et al., 2009). However, AL projection neurons were less likely to provide input to PM projection neurons than to other AL projection neurons (Figure 1h). Similarly, connections from PM to AL projection neurons were less common than between PM projecting cells (Figure 1h). There was no significant difference in distances between the somata of AL/AL and AL/PM pairs (Figure 1j, median 69.5 µm and 64.0 µm, respectively, *p* = 0.66). Although AL/PM pairs tended to be further apart than PM/PM pairs (median 64 µm and 48 µm, *p* = 0.024), this difference could not account for lack of connectivity between them. Indeed, we observed that none of the 44 AL/PM pairs closer than 48 µm were connected.

The amplitude of excitatory postsynaptic potentials between connected PM projection neurons was similar to that of AL projection neuron pairs (*p* = 0.10, two-sided Wilcoxon rank sum test; Figure 1i). Due to the rarity of connections between AL and PM projection neurons, no conclusion could be made about their connection strength. Finally, we found no statistical difference in intrinsic membrane properties of AL and PM projection neurons in L2/3 (Supplementary Figure 1) or short-term plasticity (paired pulse ratios) of postsynaptic responses of AL and PM neurons. (30 Hz, 5 ms pulse stimulation; AL to AL, 0.92 *±* 0.39 (mean *±*standard deviation), *n* = 26; PM to PM, 0.92 *±* 0.61, *n* = 11; AL to PM, 1.05 *±* 0.22, *n* = 3; PM to AL, 0.94 *±* 0.19, *n* = 4). These values are comparable to findings reported previously (Maravall et al., 2004; Oswald and Reyes, 2008).

In summary, AL and PM projection neurons in V1 have similar biophysical properties but differ in their patterns of local connectivity. They form exclusive subnetworks, preferentially making synaptic connections with neurons projecting to the same target.

### AL and PM projection neurons carry distinct visual information

How do the mutually exclusive subnetworks of AL and PM projection neurons differ in their responses to visual stimuli? To answer this question, we simultaneously imaged the activity of these cell populations in awake mice passively viewing grating stimuli. After identifying areas AL and PM using intrinsic signal imaging, we used Pseudorabies Virus (PRV)-Cre (Oyibo et al., 2014) and CTB-conjugated with AlexaFluor-647 to back-label V1 neurons projecting to these areas (Figure 2a-b). Meanwhile, we expressed GCaMP6f broadly in V1 neurons. We then imaged brain volumes spanning 320-420 µm across and 80 µm in depth 100-300 µm below cortical surface, while tracking animals’ running behaviour and eye movements (Figure 2c-d).

**Figure 2.**
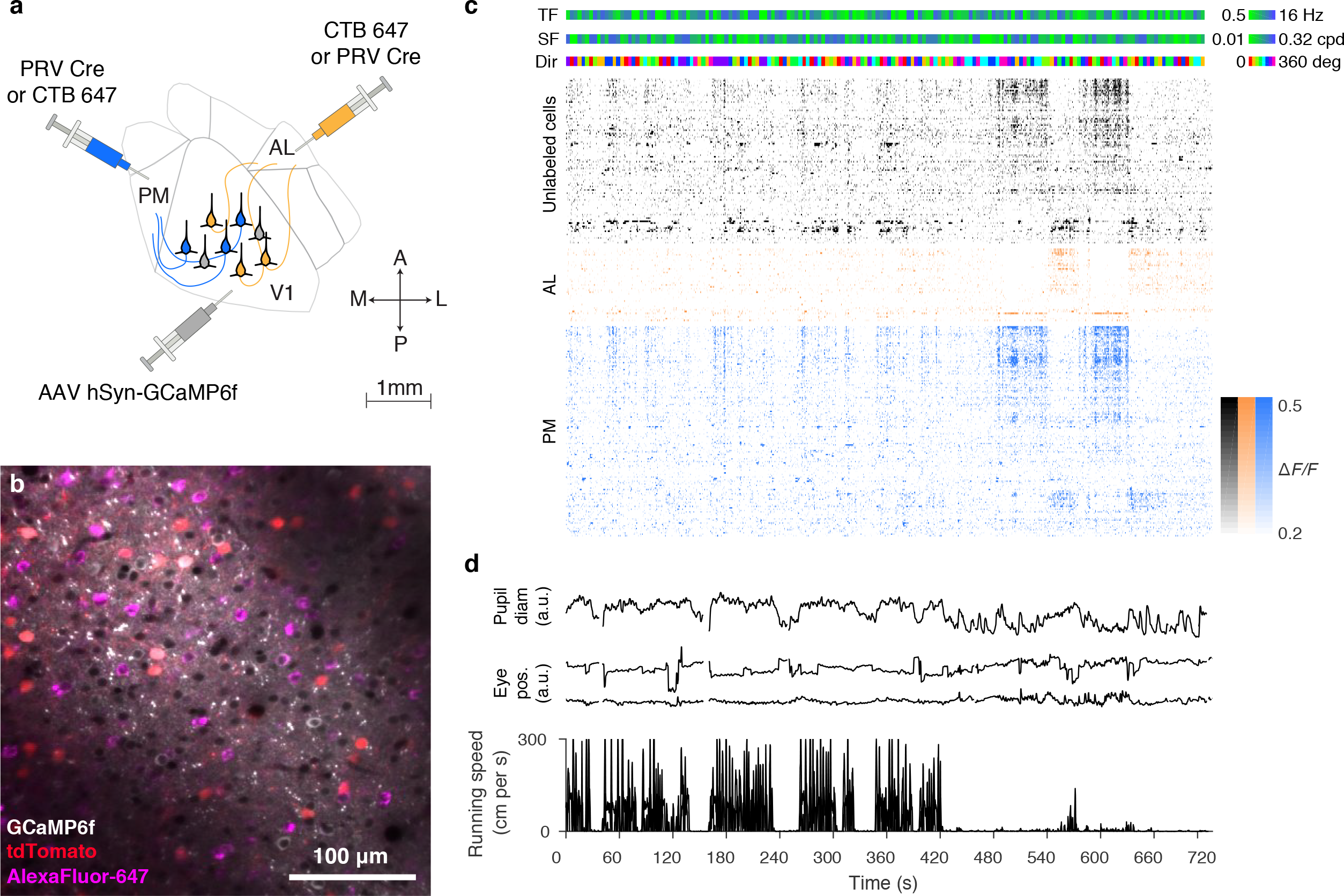
Simultaneous imaging of AL and PM projection neurons. **a**. Strategy for labelling of AL and PM projection neurons. PRV Cre was injected into either area AL or PM of Ai14 Lox-STOP-Lox tdTomato mice, while AlexaFluor-647-conjugated Choleratoxin B was injected into the other area. AAV driving expression of GCaMP6f was injected into V1. **b**. AlexaFluor-647 (magenta), tdTomato (red) fluorescence in PM and AL projection neurons, respectively. GCaMP6f fluorescence in gray. **c**. Example raster of responses of projection neurons and a random selected subset of unlabeled cells. **d.** Pupil diameter, eye position and running speed during the imaging session.

To characterize the selectivity of AL and PM projecttion neurons for different attributes of visual input, we varied the spatial and temporal frequency of the grating stimuli over 6 octaves (Figure 3a). Neurons in both sub-populations were highly heterogeneous and spanned the full range of spatial and temporal frequency preference as well as orientation and direction selectivity (Figure 3b). However, they were quantitatively different in several respects. First, PM projection neurons tended to prefer high spatial frequency stimuli (Figure 3c-e), while AL projection neurons preferred high temporal frequency and high speed stimuli (Figure 3c-d,f-g). These observations are consistent with the visual response properties of axonal boutons of V1 neurons in areas AL and PM (Glickfeld et al., 2013). In addition, reminiscent of MT projection neurons in macaque area V1 (Movshon and Newsome, 1996), cells targeting area AL tended to be more direction selective than PM projection neurons (Figure 4h).

**Figure 3.**
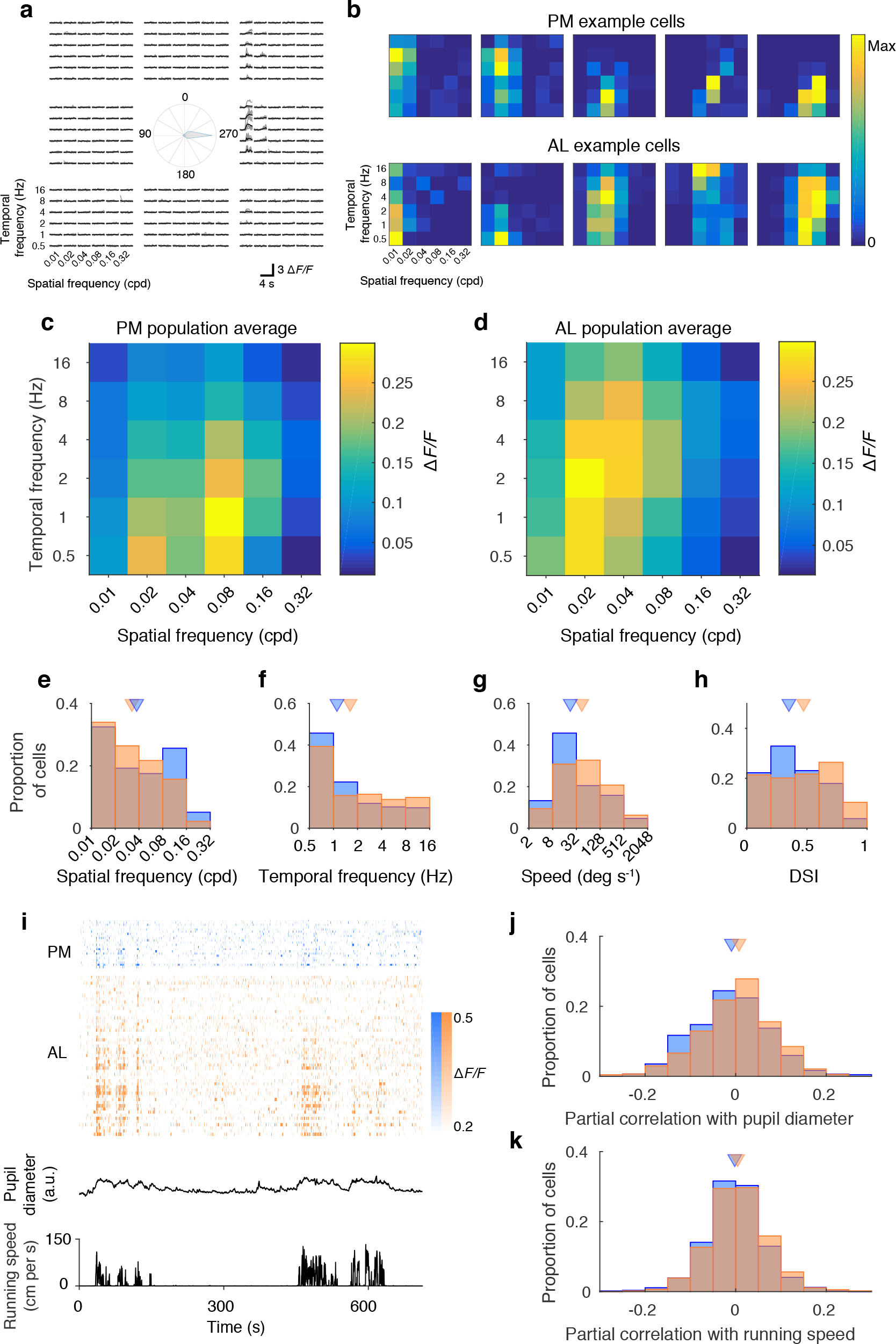
Tuning properties of AL and PM projection neurons. **a**. Responses of an example AL projection neuron to different spatial, temporal frequencies and directions. Gray-individual trials, black - mean responses. **b**. Responses of AL and PM projection neurons are diverse. Normalized responses at the preferred direction for AL and PM neurons. Examples were selected to reflect the range of their spatial frequency tuning. **c-d**. Mean △F/F responses at preferred direction for PM (c, *n* = 234 cells) and AL (d, *n* = 318 cells) projection neurons. **e-h**. Preferred spatial frequency (**e**, *p* = 0.03, Wilcoxon rank-sum test), temporal frequency (**f**, *p* = 0.0058), speed (**g**, *p* = 6.6 *×* 10^*−*5^) and direction selectivity index (**h**, *p* = 2.3 *×* 10^*−*5^) for AL (orange) and PM (blue) projection neurons. Triangles show medians. **i**. Example raster of responses of PM (blue) and AL (orange) projection neurons sorted based on correlation with pupil diameter. **j.** Responses of AL projection neurons have higher partial correlations with pupil diameter, controlling for running speed (*p* = 5.8 *×* 10^*−*5^), 1080 PM and 908 AL projection neurons). **k.** Responses of AL projection neurons have higher partial correlations with running speed, controlling for pupil diameter (*p* = 0.0061).

Behavioral state modulates the responses of neurons in mouse V1 (Neill and Stryker, 2010). To determine whether AL and PM projection neurons are differentially modulated by behavioral parameters, we examined the relationship of their responses with running speed and pupil diameter (Figure 3i). Since running speed and pupil diameter are themselves correlated with each other, to disambiguate their effects we calculated the partial correlation of single cell responses with each of these two parameters whilst controlling for the other. Responses of AL projection neurons were more strongly correlated with both pupil diameter and running speed (Figure 3j-k).

### Response connectivity correlations of AL and PM projection neurons do not explain their exclusive

In layer 2/3 of mouse V1, pyramidal neurons that show correlated responses to visual stimuli are more likely to connect to each other (Ko et al., 2011; Cossell et al., 2015). Could the similarity of responses of cell pairs projecting to the same long-range target explain the exclusive local connectivity of AL and PM projection neurons? Simultaneous recordings of AL and PM projection neurons allowed us to directly test this hypothesis. We first compared the overall similarity of activity patterns by calculating Pearson’s correlation coefficient of fluorescence traces between pairs of cells (Figure 4a). Across the population, correlations decayed with separation between cell pairs (Figure 4b; Lur et al., 2016, Ringach et al., 2016). Correlations of cell pairs projecting to the same long-range target showed the same pattern as unlabelled neurons. On the other hand, pairs of cells projecting different long-range targets were less correlated than either unlabelled cell pairs or pairs projecting to the same target. These results parallel the relative absence of local connections between AL and PM projection neurons observed in vitro (Figure 1h).

**Figure 4.**
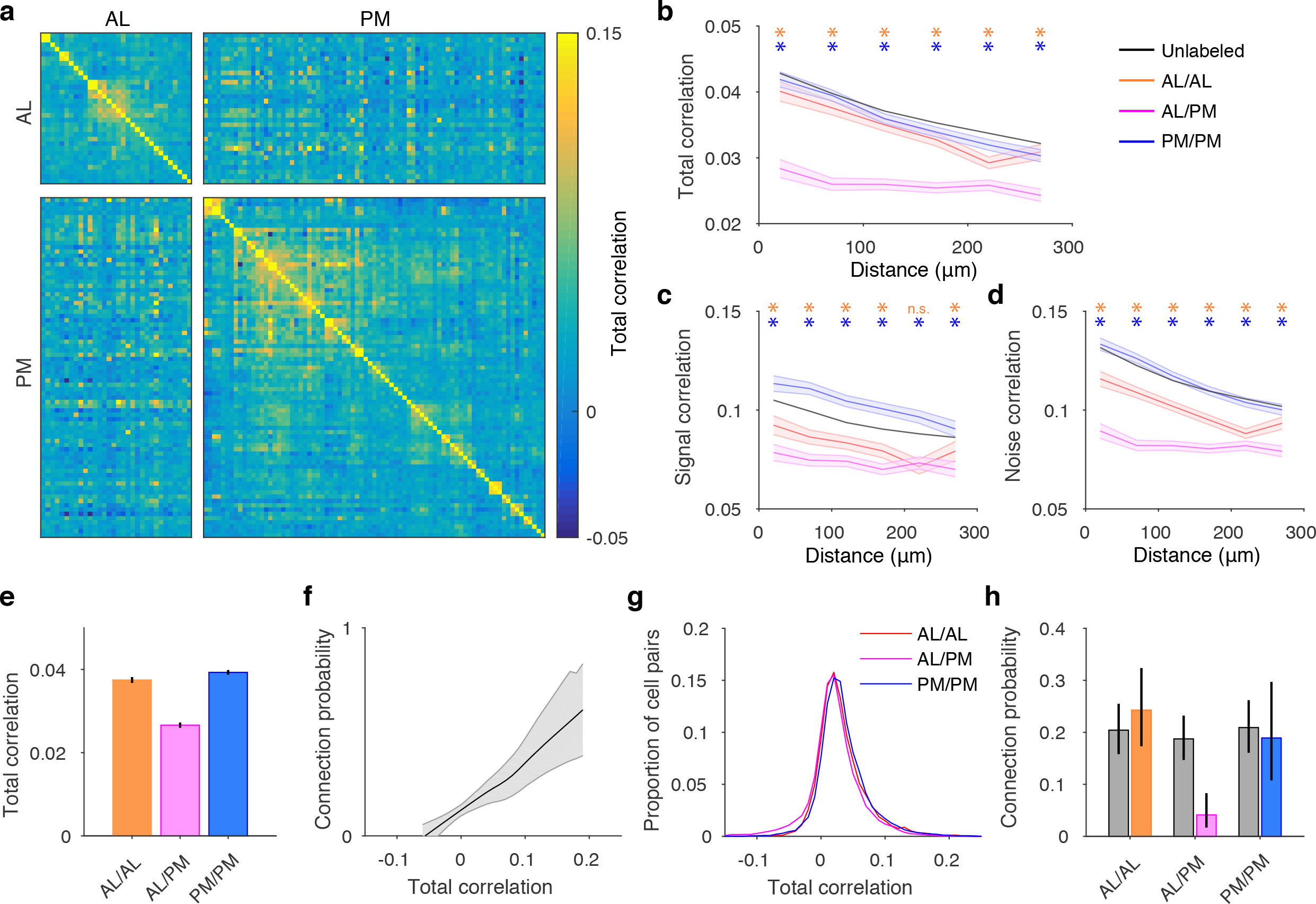
Relating correlation structure and connectivity of AL and PM projection neurons. **a.** Total correlation matrix of AL and PM projection neurons during an example session. **b.** Total correlation of unlabelled (black), AL/AL (orange), PM/PM (blue), and AL/PM (magenta) neuron pairs with respect to cortical distance. Shading denotes 95% confidence intervals estimated from standard error. Asterisks label bins, for which correlation of AL/AL (orange) or PM/PM (blue) cell pairs was significantly different from AL/PM pairs (*p <* 0.05, ranksum test). **c.** Signal correlation of unlabelled, AL/AL, PM/PM, and AL/PM neuron pairs. **d.** Noise correlation of unlabelled, AL/AL, PM/PM, and AL/PM neuron pairs. **e.** Mean total correlation for cell pairs within 150 µm. **f.** Relationship between connection probability and total correlation, based on data from (Cossell, Iacaruso et al., 2015). **g.** Distributions of total correlations for AL/AL, AL/PM and PM/PM cell pairs. **h.** Predicted (gray) and actual (color) connection probabilities for AL/AL, AL/PM and PM/PM cell pairs. All error bars are 95% confidence intervals.

To determine whether this observation could be explained by differences in tuning between AL and PM projection neurons, we compared their signal correlations, computed as the correlation coefficient of mean responses across stimulus types, and noise correlations, defined as the correlation of trial-to-trial deviation in responses from the mean (Figure 4c-d; see Methods). Although cells projecting to the same long-range target had higher signal correlations than cells targeting different areas, this difference was more pronounced for noise correlations, suggesting that it is the trial-to-trial variability in responses of projection neurons and not their visual selectivity that gives rise to the reduced correlations of cell pairs projecting to different targets.

We then used the previously reported relationship between synaptic connectivity and response correlation (Figure 4f; Ko et al., 2011; Cossell et al., 2015) to estimate the expected connectivity rates of these subpopulations, assuming they were determined solely by the similarity of their responses. We estimated the connection probability for individual cell pairs using the relationship in Figure 4f as a lookup table, and then averaged these values across each population. Although the mean correlations were lower for cell pairs projecting to different targets (Figure 4e), the correlation distributions overlapped extensively (Figure 4g). Consequently, predicted connectivity rates were similar for different cell populations (Figure 4h). Importantly, observed connectivity for cell pairs projecting to different long-range targets was significantly lower than predicted by their response correlations. This observation suggests differences in visual response selectivity alone cannot explain the lack of connections between AL and PM projection neurons and that other mechanisms must play a role in establishing the exclusive connectivity of projection neurons in mouse V1.

## Discussion

### Specific connectivity of AL and PM projection neurons

By examining the local connectivity of V1 neurons projecting to areas AL and PM, we observed that connections between neurons projecting to different targets were rare. Asymmetric connectivity has previously been described for different classes of layer 5 projection neurons (Morishima and Kawaguchi, 2006; Brown and Hestrin, 2009; Morishima et al., 2011), but has not been examined for corticocortical neurons projecting to different cortical areas.

Recent work has identified similarity of visual responses as the prime predictor of local synaptic connectivity between pyramidal cells in mouse V1 (Ko et al., 2011; Cossell et al., 2015). However, the functional properties of AL and PM projection neurons were heterogeneous and overlapped extensively. As a consequence, the expected rate of connectivity between these populations was substantially higher than we observed, indicating that projection target acts independently of response similarity in constraining local connectivity of cortical neurons. One possibility is that AL and PM projection neurons are hard-wired to avoid making connections with neurons projecting to the other target by their molecular makeup. This would imply that the molecular identity of pyramidal neurons regulates the selection of their local as well as long-range connections.

To what extent could this exclusive connectivity rule generalize to other populations of projection neurons in layer 2/3? Further work is required to determine the extent, to which the projections from V1 to other higher visual areas (Wang and Burkhalter, 2007) constitute distinct non-overlapping populations, and how these targets constrain their local connectivity. Nonetheless, estimates for the overall rate of connectivity, ranging from 10 to 20%, set an upper bound on the possible number of exclusively interconnected local subnetworks. Assuming a within population connectivity rate of 20%, as observed in our study, it is unlikely that more than two major exclusively connected subnetworks of projection neurons exist. Therefore, if similar connectivity rules apply to other populations of projection neurons, it seems likely that they would participate in same subnetworks as AL or PM projection neurons rather than forming their own subnetworks.

### Functional specialization of projections to higher visual areas

By analogy with dorsal and ventral streams of visual processing in primates, it has been proposed that projections from mouse V1 to areas AL and PM are specialized for extracting different types of information from the visual scene (Wang et al., 2011; Glickfeld el al., 2013). Although neurons projecting to areas AL and PM are biased to different preferred speeds, and temporal and spatial frequencies, both projections are highly heterogeneous. It therefore seems more probable that both of these projections relay a complete representation of the visual scene, tailored to the computational needs of different target areas.

### Correlated responses of AL and PM projection neurons

Responses of V1 pyramidal neurons projecting to the same long-range target tend to be weakly positively correlated, while cell pairs projecting to different targets have lower correlations than randomly sampled neurons. These correlations arise primarily from shared trial-to-trial variability in neuronal responses rather than differences in stimulus selectivity between different projecting populations. These observations parallel the connectivity of these cell populations and suggest that the decreased correlations between AL and PM projection neurons are a consequence of the dearth of synaptic connections between them. In essence, the lack of connections between these subpopulations appears to functionally insulate them, preventing the fluctuations in activity one population from influencing the other.

What is the impact of these correlations on sensory coding by V1 projection neuron populations? It seems reasonable to assume that higher noise correlations among neurons projecting to a particular area would limit the potential information bandwidth of that pathway. However, as recent work has demonstrated that the effects of noise correlations on information coding cannot be readily inferred from pairwise measurements (Moreno-Bote et al., 2014), this question warrants further investigation.

What is the computational benefit of the exclusive connectivity of AL and PM projection neurons, given their extensive functional overlap? The absence of recurrent connections between AL and PM projection neurons could allow top-down signals to modulate their activity independently. Reduced noise correlations between these cell populations may be a signature of this modulation. We suggest that exclusive recurrent connectivity of different populations of projection neurons may allow primary visual cortex to dynamically route information, via independent modulation of these output channels by top-down signals. Characterizing how the activity of V1 projection neurons is shaped by behavioural demands will help identify the circumstances that lead to their selective recruitment. This view extends the function of recurrent connections in layer 2/3 beyond acting purely to amplify feed-forward responses (Douglas et al., 1995; Lien & Scanziani, 2013; Cossell et al., 2015) and suggests that they also play a role in shaping the output of different projection neuron populations.

## Methods

### Animals and surgical procedures

All experiments were conducted in accordance with institutional animal welfare guidelines and licensed by the Swiss cantonal veterinary office. We used wild-type C57BL/6 mice for in vitro recordings, and Ai14 LSL-tdTomato mice for in vivo imaging experiments.

Animals were anaesthetized with 5 mg/kg midazolam, medetomidine 0.5 mg/kg, and 0.05 mg/kg fentanyl and a metal headplate was implanted exposing the skull over the right visual cortex. The skull was covered with transparent cement to prevent infection. After recovery from the implant surgery, mice were prepared for intrinsic imaging as previously described (Roth et al., 2016). Square full contrast gratings of random orientation (0.08 cycles per degree) moving at 4 Hz were presented to the left eye within a circular aperture 25^*°*^ in diameter at 0^*°*^ elevation and 60^*°*^ or 90^*°*^ azimuth on a gray screen for 2 seconds at a time with an 18 second interstimulus interval. Response maps to the apertures at either position were used to identify the location of areas AL and PM.

At least a day after the intrinsic imaging session, animals were anaesthetized with midazolam/medetomidine/fentanyl anaesthesia and retrograde tracers (Cholera Toxin B conjugated to AlexaFluor-488 and AlexaFluor-594 for in vitro experiments, Cholera Toxin B conjugated to AlexaFluor-647 and PRV-Cre for in vivo experiments) were injected into retinotopically matched locations in areas AL and PM localized using the intrinsic response maps.

For in vivo calcium imaging, AAV hSyn-GCaMP6f was injected into area V1. 7-10 days after the injections, a craniotomy 4 mm in diameter was made exposing the visual cortex under midazolam/medetomidine/fentanyl anaesthesia. A glass coverslip (4 mm diameter, 0.17 mm thickness) was implanted for chronic calcium imaging.

### Histology

To quantify the distribution of AL and PM projection neurons V1, mice injected with retrograde tracers were deeply anaesthetized with sodium pentobarbital and transcardially perfused with 4% paraformaldehyde. The brains were extracted and post-fixed overnight before being prepared for sectioning and imaging using the TissueCyte serial two-photon tomography system (Tissue-Vision). For quantification of the distribution of AL and PM neurons in Figure 1, we used one mouse injected with Cholera Toxin B AlexaFluor-647 in AL and PRV-Cre in PM and one mouse injected with Cholera Toxin B AlexaFluor-594 in AL and Cholera Toxin B AlexaFluor-488 in PM.

### In vitro whole-cell patch-clamp recording

Electrophysiological recordings in brain slices were performed in mice of both sexes, aged between postnatal 27-35. After mice were lightly anesthetized with sodium pentobarbital and transcardially perfused with a cold choline chloride based solution containing (in mM): 110 choline chloride, 25 NaHCO3, 25 D-glucose, 11.6 sodium ascorbate, 7 MgCl2, 3.1 sodium pyruvate, 2.5 KCl, 1.25 NaH2PO4, and 0.5 CaCl2 (Bureau et al., 2006) with *∼* 325 mOsm. Visual cortex slices (300-350 µm thickness) were cut coronally on a vibrating blade microtome (VT1200S, Leica Biosystems) with the same choline chloride based solution bubbled with 95% O2/5% CO2. Then, the brain slices were incubated at*∼*34^*°*^C, for 20-40 min with artificial cerebrospinal fluid (ACSF) solution containing 125 mM NaCl., 2.5 mM KCl, 1 mM MgCl2, 1.25 mM NaH2PO4, 2 mM CaCl2, 26 mM NaHCO3, 25 mM D-glucose; 315-320 mOsm adjusted by adding the amount of D-glucose, bubbled with 95% O2/5% CO2, pH 7.4. Afterwards, the brain slices were continuously maintained at room temperature before being transferred to the recording chamber.

In vitro patch-clamp recordings were performed with Scientifica multiphoton imaging microscope and a mode-locked Ti:sapphire laser (Vision-S, Coherent) at 780 nm with a Nikon 16x water-immersion objective (NA 0.8). Scanning and image acquisition were controlled by SciScan (Scientifica) and custom software written in Lab-VIEW (National Instruments). Recording pipettes were mounted on remote-controlled motorized micromanipulators (MicroStar, Scientifica). Recording pipettes were made using thick-walled filamentous borosilicate glass capillaries (G150F-4, Warner Instruments) using a horizontal puller (P-1000, Sutter Instrument) adjusted to produce pipette resistance of 7-8 MOhm with a long taper when filled with intracellular solution in 34^*°*^C ACSF. The potassium based internal solution containing 5 mM KCl, 115 mM K-gluconate, 10 mM HEPES, 4 mM Mg-ATP, 0.3 mM Na-GTP, 10 mM Na-phosphocreatine, 0.1% w/v Biocytin; osmolarity 290-295 mOsm, pH 7.2 was used. Liquid junction potentials were not corrected.

Projection neurons were targeted under visual guidance using a custom laser-scanning Dodt contrast imaging system, and simultaneous two-photon imaging for detection of AlexaFluor-488 and −594 fluorescence in labelled neurons. Whole-cell patch-clamp recordings were carried out from up to 6 cells simultaneously at *∼* 34^*°*^C, using Multiclamp 700B amplifiers (Axon Instruments) and custom-written MATLAB software (MathWorks). To test for the presence of synaptic connections, five presynaptic spikes by current injections at 30 Hz were evoked in each cell sequentially (Figure 1g), repeated 20 to 150 times, while searching for corresponding postsynaptic responses. Recordings with series resistances below 35 MOhm were included for analysis.

Connection probabilities were calculated as the number of connections detected over the number of potential connections assayed. Monosynaptic connections were identified when the membrane potential of the postsynaptic cell 7.8-12.8 ms following presynaptic stimulation (5 spikes, 30 Hz; Figure 1g) was significantly greater than during the 5 ms preceding it (Wilcoxon signed rank test *p <* 0.01; excluding the time period of electrical stimulus artefacts, e.g. connections between neurons AL2 and PM2 in Figure 1g) and the latency of the peak postsynaptic response was consistent across 5 presynaptic pulses.

Paired pulse ratios were calculated as the amplitude of the 2nd postsynaptic response divided by the amplitude of the 1st postsynaptic response. For intrinsic membrane properties of AL and PM projection neurons, step currents from −50 pA to 900 pA at 50 pA increments for 1 s were injected to determine I-V relationship (Supplementary Figure 1a-b). Detailed definitions and protocols are shown in the figure legends.

### In vivo two-photon calcium imaging

Awake mice were head-fixed and allowed to run on a styrofoam wheel. A rotary encoder monitored rotation of the wheel, while a camera recorded eye movements and pupil dynamics. Signals from the rotary encoder and eye camera were acquired using custom LabVIEW software (National Instruments). A monitor (47 cm wide) was placed 22 cm away from the eye spanning a field of view of 122^*°*^. Monitor position was adjusted such that the centre of the screen matched the preferred retinotopic location of the imaging site, as judged by two-photon fluorescence responses to grating patches flashed at different locations on the screen.

Fluorescence signals were recorded using a ThorLabs B-Scope 2 resonant scanning two-photon microscope with a Nikon 16x water-immersion objective (NA 0.8) operated using ScanImage 5.1 software. Volumes of 8 frames spanning 80 µm in depth and 320-420 µm in X/Y were acquired at 3.75 Hz using a piezo focuser (PI P-726). For identification of retrogradely imaged neurons, reference volumes of AlexaFluor-647 fluorescence were imaged at 830 nm with a 676/29 nm emission filter (Semrock), while tdTomato fluorescence was imaged at 930 nm with a 607/70 nm emission filter (Semrock). GCaMP6f fluorescence was visible at both wavelengths and was used to align the reference volumes. For functional imaging, GCaMP6f fluorescence was imaged using 930 nm excitation at 10-30 mW with a 520/40 nm emission filter (Chroma).

To prevent the light from the monitor from interfering with imaging, the monitor backlight was controlled by a custom electronic circuit and only switched on during the turn-around of the resonant X mirror (Leinweber et al., 2014). Gratings of 6 spatial frequencies, 6 temporal frequencies and 8 directions were interleaved randomly and presented without gaps for 6-8 repetitions. Each grating first remained stationary for 2.1 seconds (8 volumes), before moving for 2.1 seconds (8 volumes).

### Data analysis

Pupil diameter was estimated as the major axis of an ellipse fit to thresholded images from the eye tracking camera.

Two-photon imaging frames were registered using a phase-correlation algorithm, ROIs were selected and fluorescence time series were extracted. Projection neurons labelled with AlexaFluor-647 and tdTomato were identified manually. To correct for bleaching, fluorescence traces were high-pass filtered at 0.0019 Hz. To correct for neuropil contamination, we used the residuals of robust linear regression of each fluorescence trace against neuropil fluorescence in a 35 µm radius surrounding each cell.

To characterize the direction, spatial and temporal frequency tuning of individual cells, we fit their responses as the product of a double Gaussian in direction space and a 2D Gaussian in spatial and temporal frequency space with arbitrary orientation. If *SF_pre f_* and *T F_pre f_* are the preferred spatial and temporal frequency of the cells, and *α* is the orientation of the SF/TF tuning Gaussian, let Δ*x* and Δ*y* be defined as:

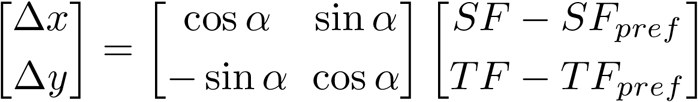

The response R is then described as:

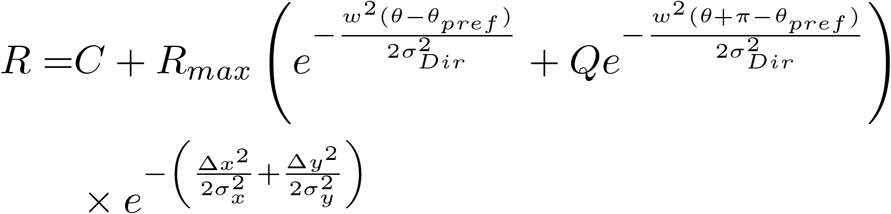

where *σ_x_* and *σ_y_* describe the tuning width of SF/TF Gaussian, *Q* is the relative magnitude of the response to the null direction (constrained between 0 and 1), *θ_pre f_* is the preferred direction, *σ_Dir_* is the width of the direction tuning, *R_max_* is the response at preferred spatial, temporal frequency, and direction, *C* is an offset, and *w*(*θ*) wraps angles onto the interval between 0 and *π*:

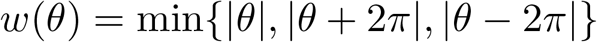

We determined the values of the fit parameters that minimized the square error of the predicted response using *lsqnonlin* in MATLAB. For the analysis of tuning properties of AL and PM projection neurons in Figure 3, we selected robustly responsive and orientation tuned neurons (*R*^2^ of the fit *>* 0.1, width of direction tuning *σ_Dir_ <* 40 degrees). However, we reached the same conclusions if we included all AL and PM projection neurons.

Total correlations were computed as the Pearson correlation coefficient of single frame fluorescence responses during the entire stimulus series. Signal correlations were computed as the Pearson correlation coefficient of the mean responses to the moving grating of each type. To calculate noise correlations, we subtracted these means from individual trial responses to the moving phase of the grating stimuli.

To estimate the relationship between total correlation and connection probability, we applied our neuropil correlation procedure to data from Cossell et al. (2015), using the mean population activity as an estimate of the neuropil signal, before calculating total correlations of responses of individual cell pairs as described above. We estimated the relationship between total correlation and connection probability using LOWESS regression. We used bootstrap resampling to derive confidence intervals for this relationship. We then estimated the predicted connection probability each population of projection neurons as the average of connection probabilities for individual pairs of neurons based on their total correlations.

## Acknowledgements

We thank Sonja Hofer for comments on the manuscript. We thank Lee Cossell for assistance with multiple whole-cell patch-clamp recording; Mohammad Hadi Saiepour for the adult mouse slicing preparation; Maxime Rio for two-photon data analysis pipeline; Rob Campbell for whole brain imaging; Georg Keller (FMI, Basel) for providing the electronic circuit design for synchronising monitor illumination to the resonant scanner. This work was supported by European Research Council (NeuroV1sion 616509 to T.D.M-F.), Swiss National Science Foundation (SNSF 31003A_169802 to T.D.M.-F.), EMBO Long-term Fellowship (ALTF 1592-2013 to P.Z.) and Marie Curie International Incoming Fellowship (PIIF-GA-2013-625740 to P.Z.).

## Author contributions

M.K., P.Z., and M.F.I. performed experiments and analysed the data. P.Z., M.K., and T.D.M-F. wrote the manuscript.

---

**Figure S1.**
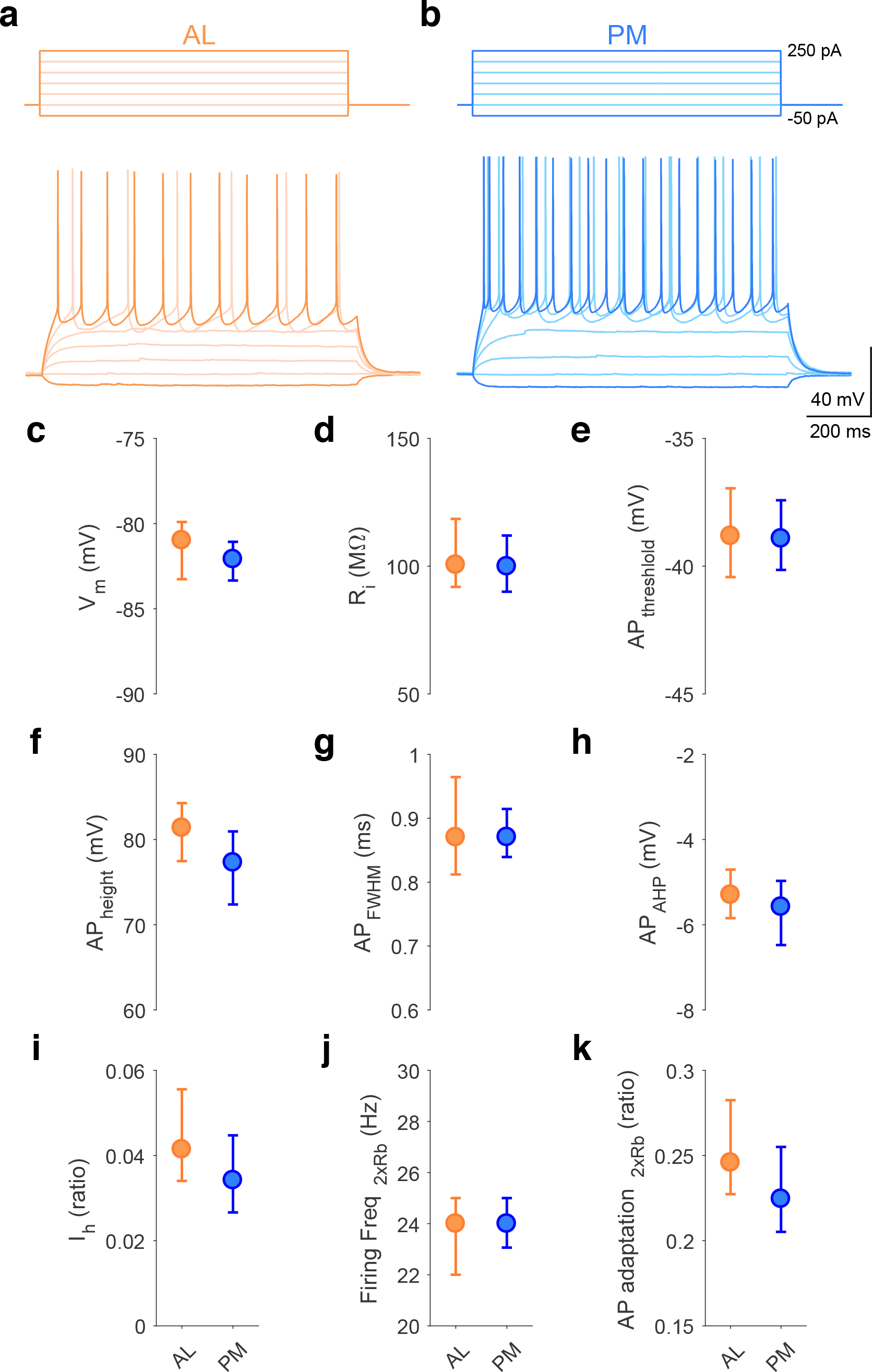
Intrinsic membrane properties of AL and PM projection neurons. **a.** Example trace of AL projection neuron (orange). Step current injection shown in upper panel and corresponding voltage response of AL projection neuron shown in lower panel. **b.** Example trace of PM projection neuron (blue). **c-k.** Circles and error bars indicate median values and 95% confidence intervals for different membrane properties. The rheobase (Rb) was measured as the lowest current injection (50 pA step with 1 s duration) leading to action potential discharge from resting potential. **c.** Resting membrane potential (V_*m*_) (AL projection neurons, n = 80; PM projection neurons, n = 88). **d.** Input resistance (R_*i*_) calculated by −25 pA current injection with 500 ms duration on the resting membrane potential (AL, n = 80; PM, n = 88). **e.** Action potential (AP) threshold (AP_*threshold*_) detected from first spike at 1x Rb (AL, n = 80; PM, n = 88) and measured from the inflection point of the minimally suprathreshold trace. **f.** AP height (AP_*height*_) detected from first spike at 1 × *Rb* (AL, n = 80; PM, n = 88) and measured by the difference between AP threshold and peak. **g.** AP width (AP_*F W HM*_) at half the peak amplitude, full width at half maximum (FWHM) (AL, n = 80; PM, n = 88) measured at the mean of threshold and peak. **h.** AP after hyperpolarization (AP_*AHP*_) defined by membrane potential at the 3 ms time point after first AP peak at 1 × *Rb* (AL, n = 80; PM, n = 87). **i.** Hyperpolarization-activated current (I_*h*_), a nonspecific cation current activated by membrane potential hyperpolarization, measured by −50 pA current injection on the resting membrane potential. The I_*h*_ potential was measured as the difference between the minimum hyperpolarized voltage from the resting membrane potential and the steady-state voltage from the hyperpolarized voltage at the end of the current step (AL, n = 79; PM, n = 88). **j.** Averaged AP firing frequency (Firing Freq_2*×Rb*_) at 2 *× Rb* (AL, n = 66; PM, n = 78). **k.** AP adaptation (AP adaptation_2*×Rb*_) defined as the the ratio of the first and the last inter-spike intervals of the spike train evoked at 2 *× Rb* (AL, n = 66; PM, n = 78).

